# Human motor neurons are rare and can be transcriptomically divided into known subtypes

**DOI:** 10.1101/2023.04.05.535689

**Authors:** Olivia Gautier, Jacob A. Blum, James Maksymetz, Derek Chen, Christoph Schweingruber, Irene Mei, Anita Hermann, David H. Hackos, Eva Hedlund, John Ravits, Aaron D. Gitler

## Abstract

We performed single-nucleus RNA-sequencing on adult human spinal cord using a neuronal nuclei enrichment strategy. We obtained transcriptomic profiles of >14,000 spinal neurons, including a small population of motor neurons that shares similarities with mouse motor neurons and can be subdivided into alpha and gamma subtypes. We sought to compare our results to those from a recent study by Yadav and colleagues, which provides a single-nucleus transcriptomic atlas of the human spinal cord. While most neuronal nuclei from both studies share similar features, our results from motor neurons differ substantially. We reanalyzed their RNA-sequencing data and provide evidence that the authors incorrectly identified cholinergic cellular debris as motor neuron nuclei in their dataset, raising doubts about their conclusions regarding motor neurons. Our findings underscore the challenges associated with transcriptionally profiling motor neurons from the spinal cord because of their rarity. We propose specific enrichment strategies and recommend important quality control measures for future transcriptional profiling studies involving human spinal cord tissue and rare cell types.

## Main Text

Motor neuron degeneration causes several devastating diseases, such as amyotrophic lateral sclerosis (ALS) and spinal muscular atrophy. Single-nucleus RNA sequencing (snRNA-seq) has emerged as a powerful approach to study the molecular complexity and cellular heterogeneity across organs and tissues, including the brain and spinal cord^1^.

We and others have previously performed snRNA-seq of the adult mouse spinal cord^2,3^. To investigate human motor neuron biology and compare it to that of mouse, we performed snRNA-seq on adult human spinal cord. Because of the rarity of motor neurons, we used an enrichment strategy. We isolated nuclei (see Methods) from the cervical spinal cord of one adult donor, stained the nuclei with an antibody for the neuron-specific marker NeuN, and used fluorescence-activated nuclei sorting (FANS) to isolate a population of ∼70% neuronal nuclei and ∼30% non-neuronal nuclei (see Methods) (Figures 1A, S1A-S1B). Previous studies have suggested that gamma motor neurons, a subtype of skeletal motor neurons, express lower levels of NeuN than other subtypes^4^—so we set gates to conservatively include even neuronal nuclei with low levels of NeuN expression. We recovered transcriptomes from 14,297 neuronal nuclei after quality control and discovered that cluster 50, which contains 40 neurons, exhibits a significant enrichment of the cholinergic marker gene *SLC5A7* as well as modest, but highly significant, enrichments of other known motor neuron marker genes such as *MNX1, ISL1*, and *CHAT* compared to other neuronal nuclei (Figure 1B-1C, Table S1)^2,5,6^. We could divide these 40 human motor neurons into two subclusters (Figure 1D), and we classified the subclusters as alpha and gamma motor neurons using known marker genes from mouse, *TPD52L1* and *VIPR2* for alpha and *PARD3B* and *CREB5* for gamma (Figure 1D-1E, Table S2)^2,3^. We also found new human-specific transcripts that will warrant future investigation (Figure 1C and 1E, Tables S1 and S2).

**Figure 1:**
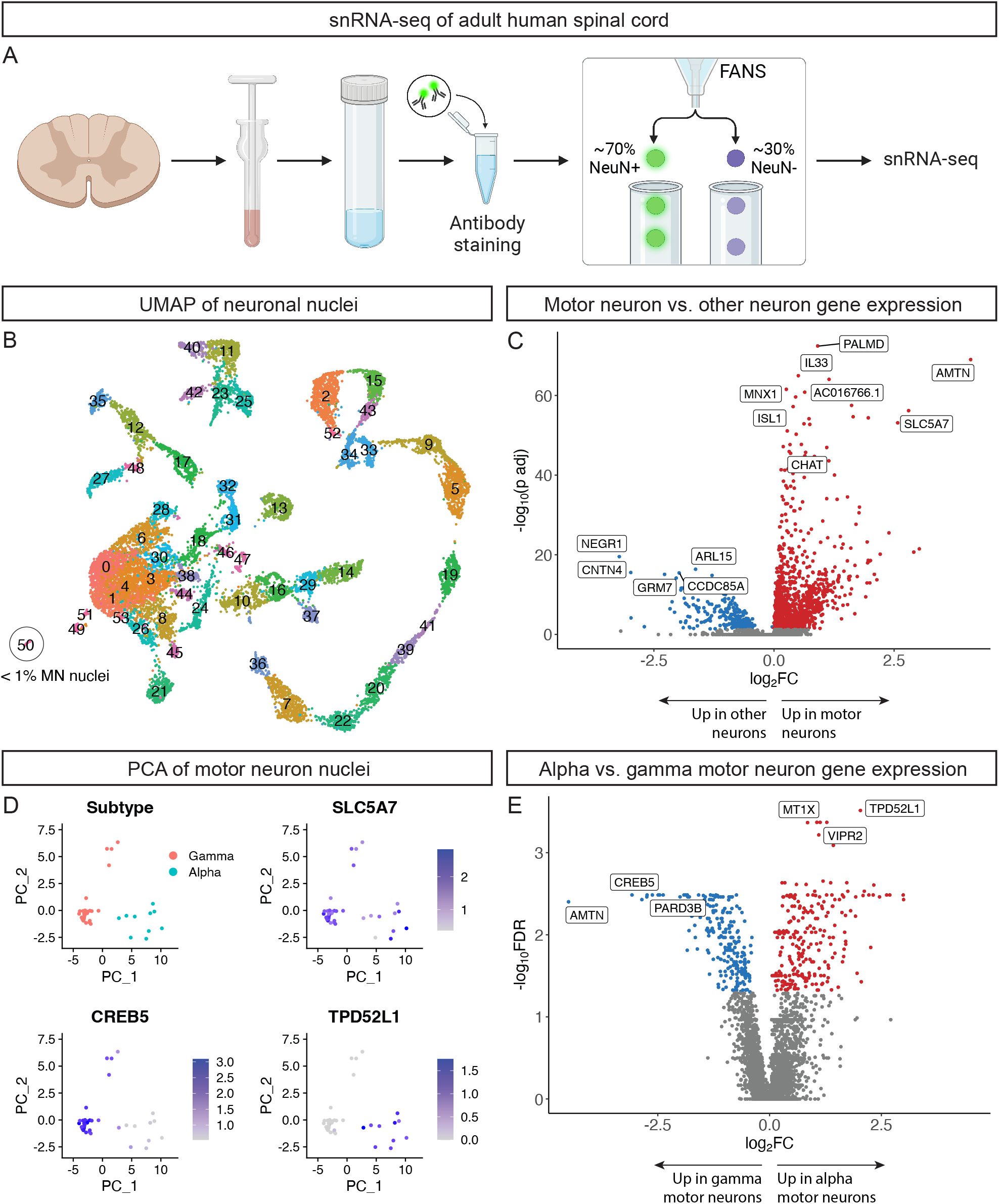
snRNA-seq reveals a small population of human skeletal motor neurons that can be divided into alpha and gamma subtypes. (A) Workflow of neuronal nuclei enrichment from human adult spinal cord followed by snRNA-seq. (B) UMAP of 14,297 neuronal nuclei. Cluster 50 corresponds to skeletal motor neurons (MNs). (C) Volcano plot showing genes significantly enriched in human motor neurons (red) or other neurons (blue) using 40 motor neurons and 500 other neurons (Wilcoxon Rank Sum test, Bonferroni-adjusted p value ≤ 0.05). (D) PCA plots showing two subclusters of motor neurons and log-normalized expression of the motor neuron marker genes *SLC5A7* (both subtypes), *CREB5* (gamma), and *TPD52L1* (alpha). (E) Volcano plot showing genes significantly enriched in alpha (red) or gamma motor neurons (blue) using 10 alpha and 30 gamma motor neurons (Wilcoxon Rank Sum test, FDR ≤ 0.05).

We sought to compare our findings to those reported by Yadav et al. who recently performed snRNA-seq on human spinal cord^7^. In addition to providing a generalized taxonomy of the human spinal cord which can be highly valuable to the research community, Yadav and colleagues focus on the transcriptional profiles of motor neurons. They report that human motor neurons are defined by the expression of cytoskeletal genes, including those related to cell size, as well as ALS genes and suggest that these features could underly the selective vulnerability of motor neurons in ALS. After cross-comparing the results with our snRNA-seq data, we identified concerns about the motor neuron transcriptional profiles reported by Yadav et al.

We first considered how Yadav et al. obtained nuclei for snRNA-seq. They isolated nuclei from the lumbar region of spinal cords from seven donor subjects. A human spinal cord contains ∼1.5-1.7 billion cells, and ∼197-222 million of these are neurons^8^. Among neurons, motor neurons are rare. Previous studies have estimated that lumbosacral human spinal cord—which is home to more than 1/3 of human skeletal motor neurons—contains ∼52-62 thousand motor neurons^9,10^. The C6 segment of the cervical spinal cord contains ∼5-6 thousand motor neurons^11^, so assuming equal distribution of motor neurons through all 8 cervical segments there should be ∼50 thousand motor neurons total in the cervical cord. Extrapolating to the entire human spinal cord, we estimate an abundance of ∼150-200 thousand total motor neurons—or approximately 0.07%-0.1% of all neurons. After various quality control steps, Yadav et al. transcriptionally profiled a total of 55,289 nuclei. Given this number of nuclei and the known composition of the spinal cord, we would expect ∼7,300 should be from neurons and only ∼5-7 of these would be from motor neurons. Depending on the homogenization and nuclei isolation protocol used, one might expect each of these numbers to vary substantially—as cell-intrinsic properties may preserve certain types of nuclei while excluding others. However, Yadav et al. report that 1,331 nuclei derive from motor neurons—around 42% of all the neuronal nuclei (1,331 out of 3,153) in their dataset (Yadav et al. Figure S11A)^7^. In other words, they claim to have recovered hundreds of times more motor neurons than expected based on cellular proportions in the spinal cord, which raises questions about the origin of these nuclei.

Another unexpected feature of the Yadav et al. motor neurons is that they express a substantially lower number of genes per nucleus compared to other neurons (Yadav et al. Figure S11C)^7^. The opposite is true in mouse where the median number of genes expressed per nucleus is ∼1.6-fold higher in motor neurons compared to other neurons^2^. Thus, we aimed to compare the number of genes and transcripts expressed per nucleus across the two datasets. In our data, the median number of genes expressed per nucleus and unique molecular identifiers (UMIs) per nucleus are ∼1.3-fold and ∼1.9-fold higher in motor neurons compared to other neurons, respectively (Figure 2A). In contrast, the median number of genes expressed per nucleus and UMIs per nucleus were < 0.2-fold and < 0.1-fold the medians of other neurons in the data from Yadav et al. (Figure 2A).

**Figure 2:**
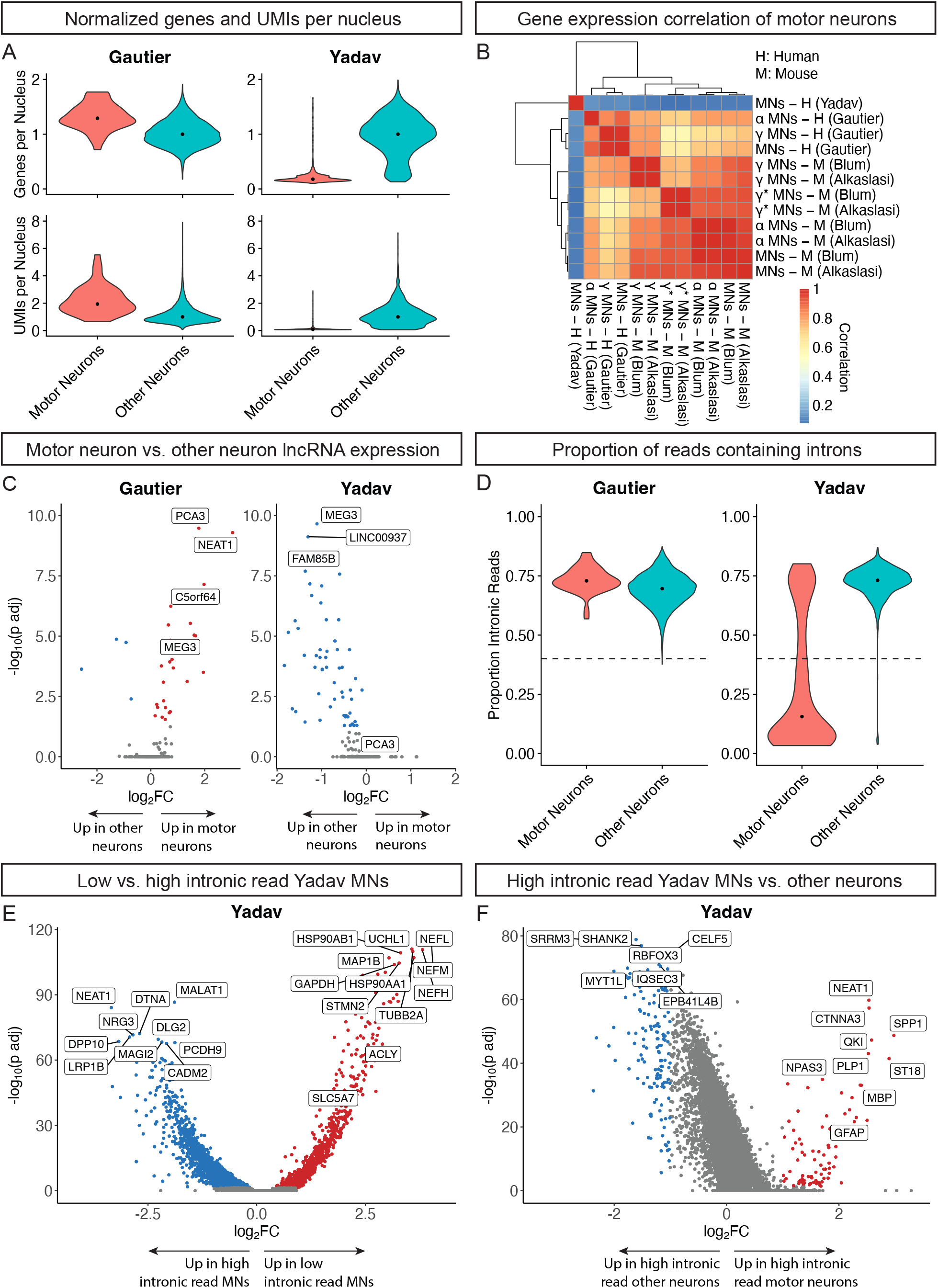
Yadav et al. motor neurons are comprised of non-nuclear, cholinergic cell debris and low-quality nuclei. (A) Violin plots showing the number of genes expressed and number of UMIs per nucleus for Gautier (left) and Yadav (right). Data are normalized to the median value of the “Other Neurons” category for each plot. Dots mark the normalized median values. Raw median values: Gautier - 8,384 genes and 42,241 UMIs for “Motor Neurons,” 6,495 genes and 21,790 UMIs for “Other Neurons”; Yadav - 936 genes and 1,432 UMIs for “Motor Neurons,” 5,249 genes and 16,809.5 UMIs for “Other Neurons.” (B) Pearson correlation of the average expression levels of mouse motor neuron marker genes for Yadav and Gautier human motor neurons as well as Blum and Alkaslasi mouse motor neurons^2,3^. (C) Volcano plot showing lncRNAs significantly enriched in human motor neurons (red) or other neurons (blue) using 40 RNA profiles per category (Wilcoxon Rank Sum test, Bonferroni-adjusted p value ≤ 0.05) for Gautier (left) and Yadav (right). (D) Violin plot showing the proportion of reads containing introns in motor neurons and other neurons for Gautier (left) and Yadav (right). Dots mark the median values, and the dashed lines are at an intronic read proportion of 0.4. (E) Volcano plot showing genes significantly enriched in low intronic read (proportion ≥ 0.4; red) or high intronic read (proportion ≥ 0.4; blue) Yadav motor neurons using 365 RNA profiles per category (Wilcoxon Rank Sum test, Bonferroni-adjusted p value ≤ 0.05). (F) Volcano plot showing genes significantly enriched in high intronic read (proportion ≥ 0.4) Yadav motor neurons (red) or other neurons (blue) using 365 RNA profiles per category (Wilcoxon Rank Sum test, Bonferroni-adjusted p value ≤ 0.05, abs(log2FC) ≥ 1).

Additionally, while the motor neurons we profiled can be subclustered into alpha and gamma subtypes (Figure 1D), this is not true of the Yadav et al. motor neurons even though they have enriched expression of two genes that are important for acetylcholine biosynthesis— *SLC5A7* and *ACLY*^7^. Further, when we compare the expression of ∼500 top motor neuron marker genes derived from mouse among the Yadav et al. motor neurons, ours, and mouse motor neurons, we find those from Yadav et al. show low correlation to other motor neurons (Figure 2B). In contrast, we found a striking correlation between mouse motor neurons and the human motor neurons we profiled (Figure 2B). Together, these observations suggested that something was awry with the population that Yadav et al. nominated as motor neurons in their snRNA-seq analysis.

Given the results above, we hypothesized that the Yadav et al. motor neuron signature may be driven by non-nuclear cell debris rather than real nuclei. One feature of nuclear transcriptomes is the presence of long non-coding RNAs (lncRNAs), which are predominantly localized to the nucleus compared to the soma. We found more lncRNAs enriched in our motor neurons compared to other neurons (Figure 2C, Table S3). In contrast, many lncRNAs are depleted from the Yadav et al. motor neurons compared to other neurons and none are enriched in motor neurons (Figure 2C, Table S4). Because splicing occurs in the nucleus, another key feature of nuclear transcriptomes is the large percentage of intronic reads from unspliced transcripts. The motor neurons we profiled have a median of ∼70% intron-containing reads, consistent with other neuronal cell populations in both studies. In contrast, the Yadav et al. motor neurons exhibit a bimodal distribution with a median of ∼16% intronic reads, consistent with a predominant subpopulation of non-nuclear cell debris (Figure 2D).

We next considered the possibility that the mode of higher intronic read motor neurons from Yadav et al. constitute real motor neurons. We split the modes at a threshold of 40% intronic reads and performed differential expression analysis on motor neurons with a high versus low percentage of intron-containing reads. We found that the motor neuron marker genes reported by Yadav et al. (Yadav et al. Figure 6A)^7^ are enriched in the population with a low percentage of intron-containing reads, including the cholinergic neuron genes *SLC5A7* and *ACLY* (Figure 2E, Table S5). We next selected Yadav et al. motor neurons and other neurons with a high percentage of intron-containing reads and performed differential expression analysis. We found that the high intronic read motor neurons from Yadav et al. do not show an enrichment of known motor neuron marker genes but instead show an enrichment of other genes including several markers for glial cells (*PLP1, MBP, GFAP*) (Figure 2F, Table S6), which may be due a higher percentage of reads coming from ambient RNA. Taken together, we believe that the motor neuron transcriptional profiles reported by Yadav et al. are not from high-quality nuclei and instead derive from non-nuclear cell debris and low-quality nuclei which have low numbers of genes expressed and UMIs detected (Figure 2A).

We found a similar population of debris in our data despite performing FANS. Cluster 22 from our initial analysis has similar features as those of Yadav et al. motor neurons, including low UMI counts, a low proportion of intron-containing reads, and enrichment of many of the motor neuron marker genes reported by Yadav et al. (Yadav Figure 6A, Figure S1A-S1C, Table S7)^7^. When comparing gene expression profiles, we found the Yadav et al. motor neurons are more similar to debris cluster 22 than they are to other neurons or our motor neurons (Figure S1D). This result suggests that similar debris will be a feature of future snRNA-seq datasets collected from adult human spinal cord and other tissue from the central nervous system.

We were still puzzled by the enrichment of cholinergic transcripts in Yadav et al. motor neurons given that these RNA profiles derive from low-quality nuclei and cell debris. To test whether there might also be non-cholinergic cell debris in the Yadav et al. dataset, we re-analyzed their data by selecting all droplets with similar characteristics as those of the motor neuron cluster— low RNA abundance (<2,000 UMIs) and high expression of neurofilament (NEFH counts>1). When applying these criteria, we were able to identify ∼2,000 droplets and subcluster them into groups defined by excitatory (*SLC17A6*), inhibitory (*SLC6A5, GAD1/GAD2*), and cholinergic (*ACLY, SLC5A7*) neurotransmitter genes (Figure S1E-S1F). We find that discrete cell debris from distinct neuronal populations are captured by this sequencing method (Figure S1G), though cholinergic events are enriched compared to debris from other neuronal types. This debris may be from small organelles known as Nissl Bodies, which are larger and much more abundant in motor neurons than in interneurons^12^, or from fragments of neurites whose membranes re-sealed. Since motor neurons have relatively large dendritic arbors and very large axon diameters^1^, it could explain why more cholinergic neuron debris is recovered. Additional investigation is needed to gain further insight into the origin of this debris.

In summary, we report transcriptional profiles from a small population of human motor neurons following enrichment of neuronal nuclei with NeuN capture and demonstrate their similarity with mouse motor neurons as well as their subdivision into alpha and gamma subtypes. We also provide evidence that the motor neuron nuclei from Yadav et al. are instead incorrectly assigned cellular debris. Our purpose in reporting these data and re-analyses is to correct the record regarding human skeletal motor neurons. They stand alone from other neurons of the spinal cord in functional and transcriptional properties and contain levels of diversity that merit further exploration. Because of their unique properties and rarity in the spinal cord, we propose that developing strategies to selectively enrich motor neuron nuclei from human (and other mammalian) tissue will empower the field to gain insights into their normal functions and the molecular etiology of motor neuron diseases.

## Acknowledgments

We acknowledge the donor whose tissue was of critical importance for this study. We also acknowledge the Chan Zuckerberg Biohub sequencing team at Stanford who helped perform the next-generation sequencing. O.G. is supported by a Stanford Graduate Fellowship. Work in A.D.G. lab is supported by NIH (grant R35NS097263). A.D.G. is a Chan Zuckerberg Biohub – San Francisco Investigator. Figure 1A was created using BioRender.com.

## Author contributions

Conceptualization, O.G., J.A.B., and A.D.G; Methodology, O.G. and J.A.B.; Investigation, O.G. and D.C.; Formal Analysis, O.G. and J.A.B.; Resources, A.H. and J.R.; Validation, J.M., C.S., I.M., D.H.H., and E.H.; Writing – Original Draft, O.G., J.A.B, and A.D.G.; Writing – Review & Editing, all authors; Visualization, O.G. and J.A.B.; Funding Acquisition, A.D.G.

## Declaration of interests

A.D.G is a scientific founder of Maze Therapeutics.

**Figure S1:**
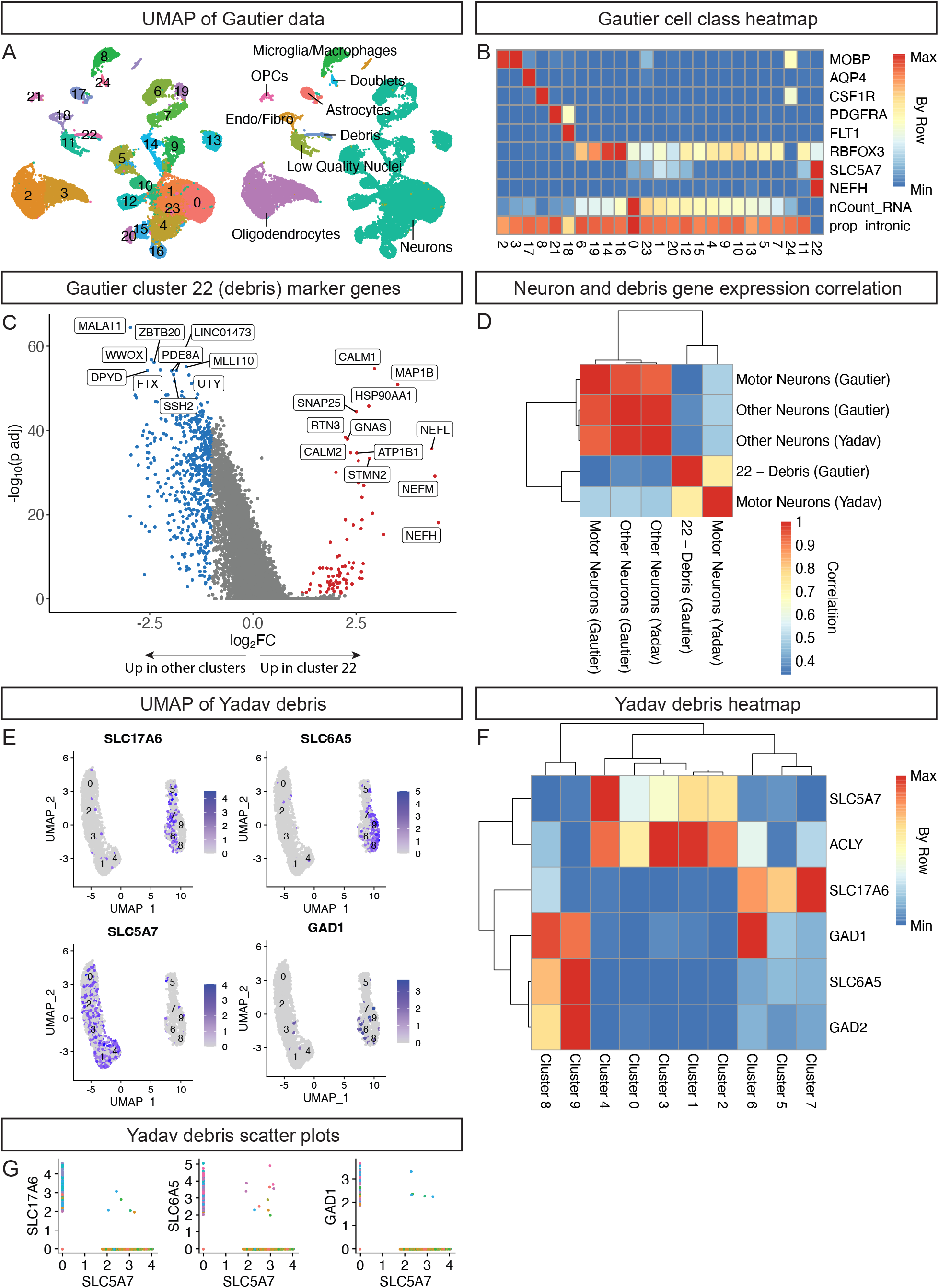
Debris from distinct neuronal subtypes is present in snRNA-seq data from human spinal cord. (A) UMAP of 21,697 RNA profiles labeled by cluster (left) and by cell class annotation (right). (B) Average expression levels of depicted genes, UMI counts (nCount_RNA), and proportion of reads containing introns (prop_intronic) for each cluster min/max-normalized by row (Transformed Value = (Value – Row Min)/(Row Max – Row Min)). These data were used to assign cell class annotations. (C) Volcano plot showing genes significantly enriched in debris cluster 22 (red) or other clusters (blue) using 214 RNA profiles per category (Wilcoxon Rank Sum test, Bonferroni-adjusted p value ≤ 0.05, abs(log2FC) ≥ 1). (D) Pearson correlation of the average expression levels of all genes detected in any category: motor neurons, other neurons, and cluster 22 debris from Gautier and motor neurons and other neurons from Yadav. (E) UMAPs of 2,005 debris profiles from a re-analysis of the Yadav data showing log-normalized expression of the neurotransmitter machinery genes *SLC17A6* (glutamatergic neurons), *SLC6A5* (glycinergic neurons), *SLC5A7* (cholinergic neurons), and *GAD1* (GABAergic neurons). (F) Average expression levels of neurotransmitter machinery genes for each Yadav debris cluster min/max-normalized by row (Transformed Value = (Value – Row Min)/(Row Max – Row Min)). (G) Scatter plots of log-normalized expression of *SLC17A6* (left), *SLC6A5* (middle), and *GAD1* (right) versus *SLC5A7* from all Yadav debris profiles.

## Methods

### RESOURCE AVAILABILITY

#### Lead contact

Further information and requests for resources should be directed to and will be fulfilled by the lead contact, Aaron Gitler.

#### Materials availability

This study did not generate new unique reagents.

#### Data and code availability

Single-cell RNA-seq data in raw and processed forms have been deposited in the Gene Expression Omnibus (GEO) with accession number GSE228778. All original code has been deposited on GitHub in the form of Jupyter Notebooks (https://github.com/neurojacob/Gautier_Blum_2023). Any additional information required to reanalyze the data reported in this paper is available from the lead contact upon request.

### EXPERIMENTAL MODEL AND SUBJECT DETAILS

Human spinal cord tissue was obtained from the UCSD ALS tissue repository, which was created following HIPAA-compliant informed consent procedures approved by an Institutional Review Board (University of California San Diego, San Diego, CA IRB# 120056). The tissue used in this study was from a 56-year-old male donor who died of an acute intracranial hemorrhage.

### METHOD DETAILS

#### Human spinal cord tissue collection

Spinal cord tissue was obtained with a postmortem interval of 2.5 hours. The tissue was dissected in the autopsy suite, flash frozen in 2-Methylbutane in a liquid nitrogen bath, and then stored at −80°C.

#### Nuclei isolation and staining

Two ∼300 mg pieces of fresh frozen cervical spinal cord were each placed in a separate 2 mL Dounce homogenizer (Sigma-Aldrich, D8938-1SET) containing 2 mL of low sucrose buffer (see Table S8; many of the buffers used are directly from or modified from Eremenko et al^13^) on ice. The tissue pieces were homogenized with 15 strokes of pestle A and 8 strokes of pestle B, and the homogenates were transferred to a 50 mL tube. The Dounce homogenizers were each rinsed with an additional 1 mL low sucrose buffer without Triton or DTT, and the solution was transferred to the same 50 mL tube. The homogenates were pelleted by centrifugation (900xg, 4°C, 5 minutes), the supernatant was discarded, and the pellets were resuspended in 4 mL 25% iodixanol solution (see Table S8). This solution was split equally into two 10 mL round-bottom centrifugation tubes (Thermo Scientific, 3118-0010PK) and then 1.5 mL 29% iodixanol solution (see Table S8) was layered underneath the solution in each tube. Nuclei were pelleted in a swinging-bucket centrifuge (3200xg, 4°C, 20 minutes), and the supernatant containing myelin and cell debris was discarded. The nuclei pellet was resuspended in 1 mL FANS buffer (see Table S8). Alexa Fluor 488-conjugated anti-NeuN antibody (Sigma-Aldrich, MAB377X) was added to the nuclei suspension at 1:1000, and the mixture was incubated for one hour (4°C, agitating). After staining, the solution was passed through a 70 µm filter (Corning, 352350). Next, the nuclei were pelleted by centrifugation (500xg, 4°C, 5 minutes) and then resuspended in 500 µL FANS buffer (see Table S8) + DAPI (2 ng/µL final concentration) prior to sorting.

#### Fluorescence-activated nuclei sorting (FANS)

Fluorescence-activated nuclei sorting was performed on a BD Biosciences FACSAria II flow cytometer using the 85 µm nozzle. Briefly, single nuclei were gated using side scatter (SSC) and DAPI measurements to ensure that debris and multiplets were gated out. After this initial gating, NeuN+ nuclei were identified on a NeuN/SSC scatterplot. These NeuN+ nuclei formed a separate mode from NeuN-non-neuronal nuclei, enabling a simple gating strategy. Approximately 100,000 nuclei were sorted at a ratio of 70% NeuN+ and 30% NeuN-into a single well of a PCR plate (Sigma-Aldrich, EP0030129504-25EA) containing 50 µL FANS buffer (see Table S8). This nuclei mixture was pelleted (500xg, 4°C) for 10 minutes, and the supernatant was aspirated until ∼15 µL remained. The nuclei were resuspended in the remaining solution, and 1 µL was used to quantify the nuclei concentration with a hemocytometer before loading nuclei into droplets according to 10x Genomics criteria (two reactions each with a targeted nuclei recovery of 10,000).

#### Droplet-based snRNA-seq

For droplet-based snRNA-seq, libraries were prepared using the Chromium Single Cell 3′ Reagent Kit v3.1 according to the manufacturer’s protocol (10x Genomics). The generated snRNA-seq libraries were sequenced using the Illumina NextSeq 2000 P3 platform with the run parameters specified in the 10x Genomics protocol. The libraries were sequenced to an average read depth of ∼59,000 reads per nucleus.

#### Analysis of snRNA-seq data

Sequencing reads were demultiplexed and aligned to the GRCh38-2020-A human reference transcriptome (10x Genomics) using Cell Ranger software (v6.1.2; 10x Genomics). For alignment, intronic reads were included in the Cell Ranger count pipeline using the “include-introns” flag. The Cell Ranger aggr pipeline was used to normalize the libraries to the same sequencing depth and generate a combined count matrix. The filtered count matrix was used, which excludes putative empty droplets.

#### Normalization and clustering

Subsequent analyses were performed in R (v4.1.1) using the Seurat package (v4.0.4). A Seurat object was created using the filtered count matrix. Mitochondrial reads were removed and then feature counts for each putative nucleus were normalized (normalization.method = “LogNormalize” in Seurat NormalizeData function). Principal component analysis (PCA) was performed on scaled data with 33,000 variable features, and the top 20 components were used for clustering and uniform manifold approximation and projection (UMAP) visualization. For clustering, the resolution parameter was set to 0.5.

The clusters were manually annotated using the average expression of known marker genes, number of UMIs, and proportion of sequencing reads containing introns per cluster (Figure S1A-S1B). To calculate the proportion of sequencing reads that contain introns, the Cell Ranger count pipeline was run with and without the “include-introns” flag. The proportion of intronic reads was calculated per putative nucleus by first subtracting the number of UMIs (nCount_RNA) excluding intronic reads from nCount_RNA including intronic reads and then dividing by nCount_RNA including intronic reads. The proportion of intronic reads was added to the meta data of the Seurat object for each putative nucleus.

#### Subclustering of neurons

Clusters labeled as neurons from Figure S1A-S1B were used for subclustering. PCA was performed on scaled data with 33,000 newly identified variable features, and the top 50 components were used for clustering and UMAP visualization. For clustering, the resolution parameter was set to 0.5. A cluster of neuronal/non-neuronal doublets was identified by high expression of the glial genes *MBP, GFAP*, and *CSF1R*. The doublet cluster was removed, and the above process was repeated but with the clustering resolution parameter set to 2.5 to produce the UMAP of neuronal nuclei in Figure 1B. There was no further attempt to remove remaining doublets.

#### Subclustering of motor neurons

Cluster 50 from Figure 1B was used for subclustering. PCA was performed on scaled data with 40 variable features, and the top 2 components were used for clustering (resolution parameter set to 0.4) and visualization. The two resulting subclusters were manually annotated as alpha and gamma motor neurons using the expression of known marker genes from mouse^2^.

#### Normalization of Yadav et al. data

The Seurat object from Yadav et al.^7^ was obtained from the authors. The count data were log-normalized (normalization.method = “LogNormalize” in Seurat NormalizeData function).

#### Mapping mouse genes to human orthologs

Mouse spinal cord snRNA-seq data from Blum et al.^2^ were downloaded from http://spinalcordatlas.org/assets/downloads/all.exps.integrated.RDS. Top mouse skeletal motor neuron marker genes from Blum et al. 2021^2^ were identified using the FindMarkers function on skeletal motor neurons versus all other cell types (logfc.threshold = 0.25, max.cells.per.ident = 500, only.pos = TRUE). An integrated data set from Blum et al^2^. and Alkaslasi et al.^3^ was downloaded from http://spinalcordatlas.org/assets/downloads/lepichon_gitler_integrated.RDS. Mouse gene names were converted to corresponding human gene names using the biomaRt package (v2.50.3; host = “https://dec2021.archive.ensembl.org/“ in useMart function). Only marker genes with a 1:1 mouse ortholog to human ortholog relationship were used for Figure 2B (see original code).

#### Differential lncRNA expression analysis

Human lncRNA gene names were obtained using the biomaRt package (v2.50.3; biomart = “ENSEMBL_MART_ENSEMBL”, dataset = “hsapiens_gene_ensembl” in the useMart function; attributes = c(“hgnc_symbol”), filters = “biotype”, value = “lncRNA” in the getBM function). Only lncRNAs expressed in at least 10% of motor neurons or other neurons were used for each plot in Figure 2C and in the corresponding tables (min.pct = 0.1 in FindMarkers function). See figure legend for more details.

#### Intronic read analysis

10x Genomics BAM files from the seven donors from Yadav et al.^7^ were downloaded from the Sequence Read Archive (SRA; SRP349799). The BAM files were converted to FASTQ files using the Cell Ranger bamtofastq tool. The proportion of sequencing reads that contain introns was calculated as described above using the raw feature matrices. The raw data for Tsai1A could not be found, and thus, the data from Tsai1A were excluded from the analyses in Figure 2E-2F.

#### Debris droplet analysis

All original code is available on <u>Github</u>. Briefly, H5 matrices derived from seven donors sequenced by Yadav et al. were downloaded from GEO (accession number: GSM5723843). Sequenced 10x droplets were filtered based on characteristic features of the Yadav et al. motor neuron cluster—the presence of *NEFH* transcripts (*NEFH*>1) and low UMI counts (100<UMI<2,000). All droplet transcriptomes were log-normalized and Seurat was used to FindVariableFeatures (n=2,000) and then integrate across samples using default parameters. Clusters that did not share characteristic features of Yadav motor neurons (ex: high *NEAT1* expression and low *NEFM* expression) were removed, and droplets were re-clustered. This process of iterative cluster removal was repeated twice. All other analysis was performed as above.

### QUANTIFICATION AND STATISTICAL ANALYSIS

Statistical tests and parameters are reported in the figure legends. Statistical analyses were performed using the Seurat package (v4.0.4) in R (v4.1.1).

## Supplemental information

- Figure S1: Debris from distinct neuronal subtypes is present in snRNA-seq data from human spinal cord
- Table S1: Differential expression analysis – Gautier MNs vs. other neurons, related to Figure 1
- Table S2: Differential expression analysis – Gautier alpha vs. gamma MNs, related to Figure 1
- Table S3: Differential lncRNA expression analysis – Gautier MNs vs. other neurons, related to Figure 2
- Table S4: Differential lncRNA expression analysis – Yadav MNs vs. other neurons, related to Figure 2
- Table S5: Differential expression analysis – Yadav low vs. high intronic read MNs, related to Figure 2
- Table S6: Differential expression analysis – Yadav high intronic read MNs vs. other neurons, related to Figure 2
- Table S7: Differential expression analysis – Gautier debris cluster 22 vs. other clusters, related to Figure S1
- Table S8: Buffers used in nuclei isolation, staining, and FANS, related to STAR Methods

